# PINK1 Regulates Dopamine and Lipids at Mitochondria to Maintain Synapses and Neuronal Function

**DOI:** 10.1101/814343

**Authors:** Christine Bus, Sven Geisler, Marita Feldkaemper, Hector Flores-Romero, Anna Schaedler, Katharina Zittlau, Maria Zarani, Betül Uysal, Nicolas Casadei, Petra Fallier-Becker, Lisa Schwarz, Jos F. Brouwers, Henner Koch, Aslihan Ugun-Klusek, Klaudia Maruszczak, Daniela M. Vogt Weisenhorn, Wolfgang Wurst, Benjamin Schmidt, Gerard Martens, Britta Brügger, Doron Rapaport, Ana Garcia, Boris Macek, Rejko Krüger, Thomas Gasser, Philipp Kahle, Julia C. Fitzgerald

## Abstract

Mitochondrial dysfunction contributes to the pathogenesis of Parkinson’s disease but it is not clear why inherent mitochondrial defects lead specifically to the death of dopaminergic neurons of the mid brain. PINK1 is mitochondrial kinase and *PINK1* mutations cause early onset Parkinson’s disease.

We found that in neuronal progenitors, PINK1 regulates mitochondrial morphology, mitochondrial contact to the endoplasmic reticulum (ER) and the phosphorylation of Miro1. A compensatory metabolic shift towards lipid synthesis provides mitochondria with the components needed for membrane renewal and oxidative phosphorylation, maintaining the mitochondrial network once mature.

Cholesterol is increased by loss of PINK1, promoting overall membrane rigidity. This alters the distribution of phosphorylated DAT at synapses and impairs dopamine uptake. PINK1 is required for the phosphorylation of tyrosine hydroxylase at Ser19, dopamine and calcium homeostasis and dopaminergic pacemaking.

We suggest a novel mechanism for PINK1 pathogenicity in Parkinson’s disease in addition to but not exclusive of mitophagy. We also provide a basis for potential therapeutics by showing that low doses of the cholesterol depleting drug ß-cyclodextrin reverse PINK1-specific phenotypes.

## Introduction

*PINK1* mutations are the second most frequent cause of early-onset PD and *PINK1* variants have been found in sporadic PD but also in healthy controls^1, 2^. PINK1 PD has an early occurrence of L-DOPA-associated dyskinesia, slow disease progression and absence of cognitive impairment^3, 4^ and affective and psychotic symptoms are frequently part of the clinical presentation^5^.

To unravel the molecular function of PINK1, Knockouts (KOs) of the PINK1 gene in mice, drosophila, zebrafish and cancer cells have been generated indicating that PINK1 deficiency causes mitochondrial dysfunction, altered mitochondrial morphology, disturbed mitochondrial quality control and iron and calcium toxicity^6–15^. In PINK1 *in vivo* models, neurodegeneration has only been observed in zebrafish^16, 17^ with dopamine release and synaptic plasticity being affected in the striatum of mice^18^.

PINK1 acts together with another PD gene product, the E3-ubiquitin ligase Parkin ^19, 20^ to regulate mitophagy^21–23^. Upon loss of mitochondrial membrane potential or accumulation of misfolded proteins, PINK1 stabilizes on the outer mitochondrial membrane^23–25^. PINK1 then phosphorylates ubiquitin at Ser65 to activate Parkin activity^26–28^. Parkin ubiquitinylates mitochondrial outer membrane proteins such as Mitofusins^29^ and Miro^30^ which is also targeted for phosphorylation^31^. The buildup of ubiquitin chains on the outer mitochondrial membrane acts as a signal for the recruitment of autophagy receptors^21, 32, 33^ important for mitochondrial clustering^34, 35^. Some *in vivo* studies have shown that PINK1 is not required for basal mitophagy in neurons^36, 37^ or in human platelets^38^ but the topic is controversial (for an extensive review see^39, 40^) and therefore much more work is needed.

The pathogenic action of PINK1 mutations could be cell type specific. PINK1 is highly expressed in the brain but it is also expressed throughout the rest of the body and has been associated with disease mechanisms in several tissues^41, 42^ including the progression of some cancers^14, 43^. PINK1 is reportedly expressed predominantly in neurons^44^ but has been found highly expressed in myelinating oligodendrocytes in the human cortex^45^. First studies in human PINK1 patient and stem cell-derived neurons described defective Parkin recruitment to mitochondria and increased mitochondrial biogenesis^46^.

Although autophagosomes are generated at ER-contact sites^47^ PINK1 may play important roles in addition to mitophagy at mitochondrial-ER contact sites and in ER stress^48–50^. Proteins such as Miro1, VDAC and mitofusins have been strongly linked to ER and PINK1 function relevant to PD^21, 51–54^. Mitochondrial-ER contact sites are regulatory hubs for calcium, iron, lipids and dopamine. PINK1 loss of function causes calcium defects in neurons^9, 17, 55^ and iron toxicity^6, 56–58^. A PINK1-specific RNAi screen identified lipid regulation as a key factor in PINK1 mechanism of action^59^, The lipid cardiolipin externalizes on the mitochondrial outer membrane as a signal for mitophagy^60^ and promotes the transfer of electrons to complex I to protect against PINK1 loss of function^61^. PINK1 also regulates the enzyme cardiolipin synthase^62^. Dopamine synthesis and degradation is primarily regulated at the mitochondrial outer membrane and ER because of the localization of monoamine oxidase (MAO) and tyrosine hydroxylase (TH), which require ubiquitination^63, 64^ and phosphorylation (for a review see^65^) for their activation respectively.

Recently PINK1 has been shown to mediate STING-induced inflammation^66^ and mitophagy associated with innate immunity via mitochondrial release of damage factors^67^. Furthermore, PINK1 KO mice are vulnerable to infection which induces PD-like symptoms^68^. Cholesterol, fatty acids and other modified lipids are well known to activate inflammatory pathway and therefore more work is needed to confirm PINK1s mechanisms relevant to PD.

We find that PINK1 plays an important regulatory role in fine tuning mitochondrial contacts, lipid synthesis, membrane fluidity, calcium homeostasis and dopamine metabolism targets including Miro1, TH and mitochondrial Aconitase at the mitochondria and set out a mechanism involving a metabolic shift towards cholesterol synthesis, impacting neurotransmitter uptake and dopamine metabolism, providing a link between PINK1-mitochondrial function and dopaminergic vulnerability in PD.

## Results

### Absence of PINK1 in hDANs inhibits ionophore-induced mitophagy but does not affect mitochondrial morphology

We generated iPSCs from a healthy female individual that we previously characterised^69^ and then introduced a homozygous deletion of *PINK1* using TALEN directed to Exon 1 (Figure 1A). 28-35 day old human mid-brain specific dopaminergic neurons (hDANs) were derived via neural precursor cells (NPCs) using a modified version of a differentiation protocol we previously described^69^. We selected two clonal lines that were correctly edited with no random integration where no *PINK1* transcripts upstream or downstream of the gene edit could be detected (Figure S1A). The derived hDAN cultures varied in their neuronal subtype markers between independent differentiations. hDANs presented staining levels for mature neuronal markers (Figure S1B). The neuronal marker MAP2 was abundant but the percentage of hDAN cultures positive for the dopaminergic neuron marker; forkhead-box-protein A2 (FOXA2), Tyrosine Hydroxylase (TH) and Dopamine Transporter (DAT) differed (Figure 1B). The average percentage of TH positive neurons in all hDAN cultures was approximately 25% (Figure 1B). Similar to the immunocytochemical data, TH gene expression was increased different between hDAN genotypes.

**Figure. 1.**
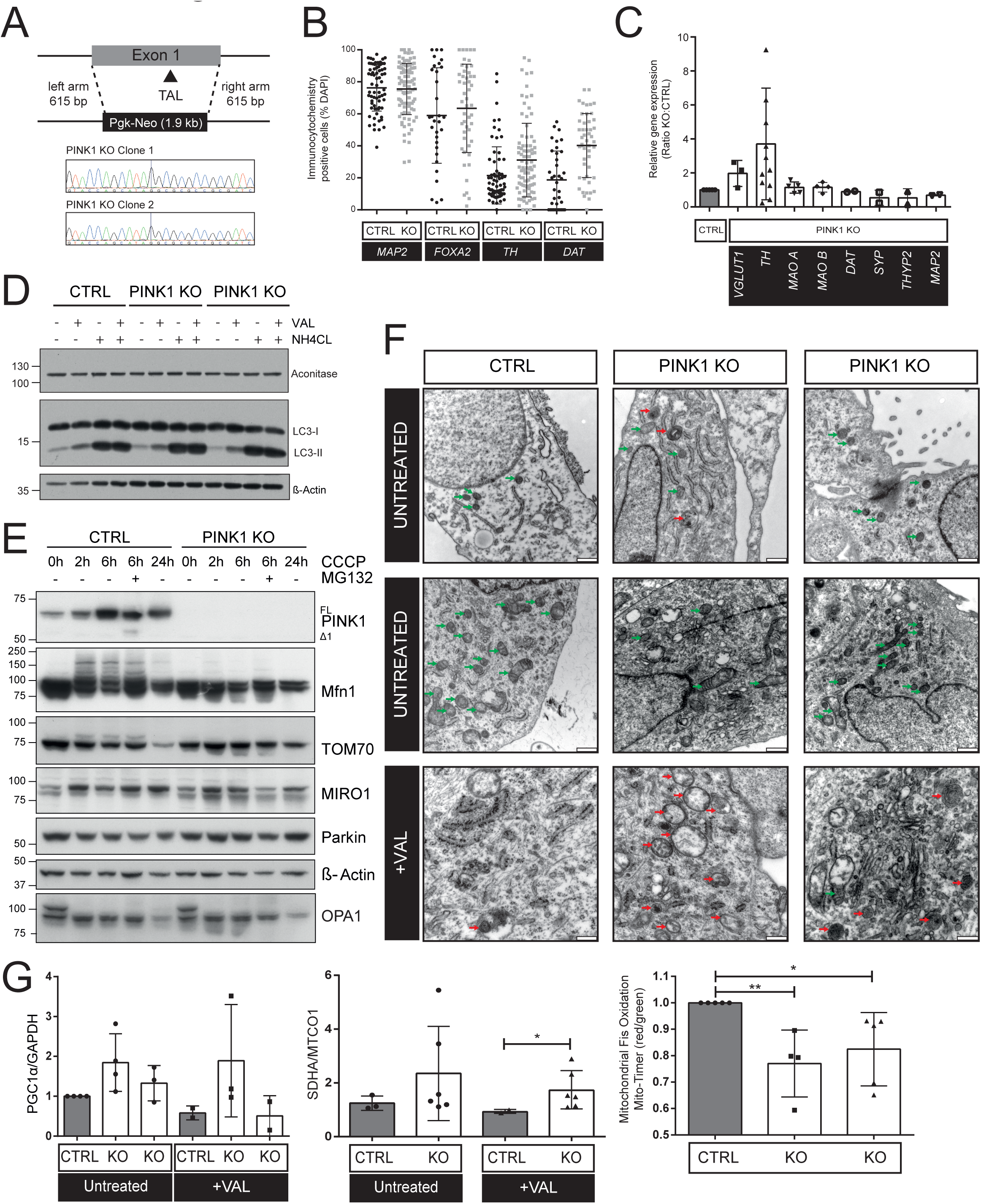
**A)** Scheme showing removal of PINK1 exon 1 (homozygous) using TALEN gene editing in healthy iPSCs. Sequence confirmation of two clonal lines. **B)** Percent of hDANs in a field of view positive for MAP2, FOXA2, TH and DAT using immunocytochemistry. Each point on the graph is a technical replicate from four independent differentiations (ndiff=4). **C)** Relative gene expression of neuronal markers in PINK1 KO hDANs compared to the healthy control. Each point on the graph is the mean average gene expression from RNA isolated from an independent hDAN differentiation. **D)** Western blots showing autophagic flux (LC3-II accumulation c.a LC3-I, Aconitase and β-actin) in untreated hDANs and those stimulated by valinomycin (Val, 1µM, 24h) or ammonium chloride (NH4Cl, 50 mM, 1h) representative blots (ndiff=3). **E)** Western blots of PINK1-Parkin pathway proteins in hDANs untreated or following 10µM CCCP treatment for 0,2,6 and 24h −/+ proteasome inhibition (MG132, 10µM, 6h). Representative blots (ndiff=3). **F)** Electron microscope images of hDANs untreated or treated with valinomycin (Val, 1µM, 24h). Representative images (ndiff=3). **G)** Quantification of Western blots for PGC1a (as a ratio to cytosolic marker GAPDH, left panel) and nuclear encoded SDH (as a ratio to mitochondrially expressed MTCO1, middle panel) (ndiff=3). Quantification of mitochondrial renewal measured by the amount of expressed Fis MitoTimer construct (red) as a ratio to oxidised Fis (green) where oxidation represents time present (right panel). Quantification of live cell microscopy images was performed blinded using Imaris software (ndiff=4).

PINK1 KO has no impact on autophagic flux in hDANs (Figure 1D), but markers of classical, CCCP-induced mitophagy such as ubiquitination of mitofusin and degradation of TOM70 were strongly reduced following a time course of 10µM CCCP treatment (Figure 1E). Steady state levels of Miro1 and OPA1 remain largely unchanged by PINK1 KO (Figure 1E). We employed the ionophore Valinomycin to promote mitophagy over a longer 24h time course to be able to fix cells (with less variability) for electron microscopy (EM). EM images from independent differentiations concluded there to be no notable differences of any organelles including mitochondrial size, shape, or cristae in untreated PINK1 KO hDANs (Figure 1F). Following mitophagy induction with Valinomycin, mitochondria (green arrows) were scarce in control hDANs, whilst mitochondria remain visible in PINK1 KO hDANs receiving the same treatment (Figure 1F). The appearance of structures resembling mitochondrial derived vesicles (MDVs) appear to increase in PINK1 KO hDANs treated with the ionophore (red arrow). Inhibited mitophagy by PINK1 KO in combination with ionophores leads to increased PGC1α levels, mitochondrial biogenesis (measured by the ratio of nuclear encoded SDHA: mitochondrial encoded MTCO1 proteins) and mitochondrial renewal (measured by the ratio of oxidized: non-oxidised mitochondrially expressed MitoTimer protein on the mitochondrial outer membrane) compared to control hDANs receiving the same treatment (Figure 1G) suggestive of compensatory PINK1-independent mitophagy and/or mitochondrial renewal.

### PINK1 knockout promotes metabolic rewiring towards lipid synthesis in hDANs

We next looked at whether PINK1 is important for mitochondrial function. Mitochondrial oxygen consumption and spare respiratory capacity are not significantly affected although PINK1 KO hDANs consume more oxygen when uncoupled and prefer glycolysis under minimal respiration conditions than their control hDANs (Figure 2A). PINK1 KO hDANs prefer not to use fatty acids for oxygen consumption (Figure 2A). These data suggest that PINK1 KO hDANs do not well metabolize acetyl CoA from glucose nor fatty acids. We asked whether complex I (CI) dysfunction may explain the shift and CI dysfunction has previously been associated with PINK1-PD^61, 70, 71^. CI enzyme activity in hDANs is not significantly affected by PINK1 KO (Figure 2B). However, since we observed variable citrate synthase enzyme activity (Figure 2C) which is used for normalization of mitochondrial mass for CI activity assays, a possible role of complex I in the PINK1 mechanism cannot be ruled out. We also found no measurable differences in the amount of active CI enzyme pulled down from untreated control or PINK1 KO hDANs (Figure S1C) and active CI (which is depleted in control hDANs following 24h valinomycin treatment) is unaffected by valinomycin in the absence of PINK1 (Figure S1C).

**Figure 2.**
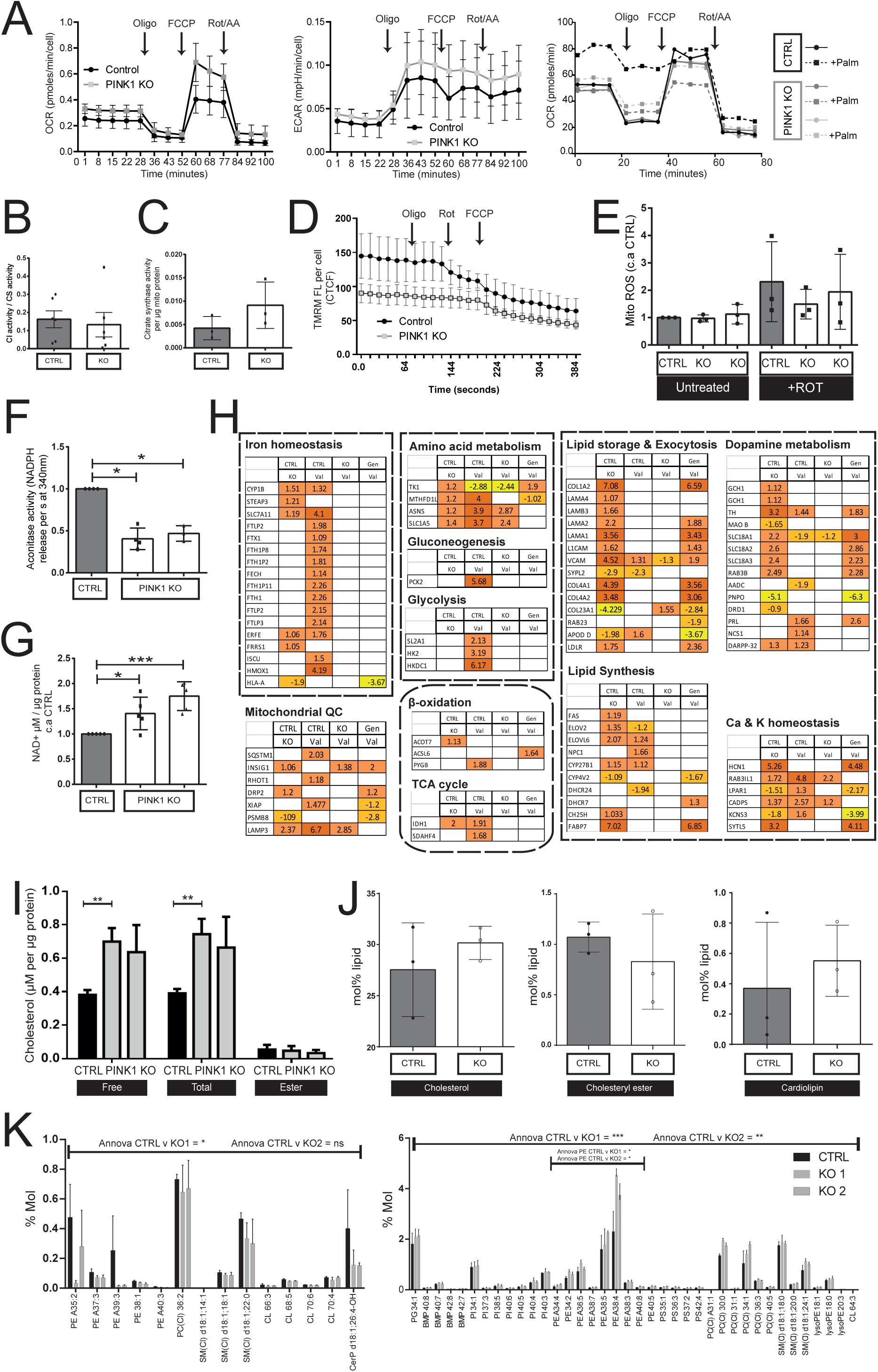
**A)** Respiratory analyses of oxygen consumption (left panel) and extracellular acidification rate (middle panel) during mitochondrial stress for hDANs where glucose and pyruvate is given and oxygen consumption (right panel) during mitochondrial stress in hDANs where carnitine and glucose is given with or without the exogenous fatty acid palmitate (Palm). **B)** NADH dehydrogenase (complex I) enzyme activity in enriched mitochondrial fractions from neurons where NADH and ubiquinone is given, NADH depletion rate in the presence of rotenone is accounted for and then normalised to citrate synthase activity as a marker of mitochondrial mass (n=diff5). **C)** Citrate synthase enzyme activity in enriched mitochondrial fractions from hDANs and normalised to total protein concentration of the mitochondrial preparation (n=diff3). **D)** Mitochondrial membrane potential measured by TMRM fluorescence quantified from live cell images per cell over a time course of where oligomycin, Rotenone and then FCCP are sequentially added (n=diff3). **E)** Mitochondrial reactive oxygen species (Mito ROS) measured using MitoSox fluorescence by flow cytometry with or without acute rotenone treatment to generate ROS. Mean fluorescence values were normalised to the background fluorescence of the same hDANs without the dye and normalised to the untreated control (n=diff3). **F)** Aconitase enzyme activity in hDANs normalised to cell number and healthy control (n=diff3). **G)** NAD^+^ concentration of hDANs normalised to total protein and to the healthy control (n=diff5). **H)** Pathway analysis of RNAseq data from hDANs where each pathway shows log2 gene expression level comparing genotype in untreated hDANs (CTRL v KO), effect of 24h ionophore treatment on control hDANs (CTRL v Val), effect of 24h ionophore treatment on PINK1 KO hDANs (KO v Val) and genotype plus treatment change (Gen v Val). **I)** Free cholesterol, total cholesterol and cholesterol ester levels in hDANs normalised to total protein levels (n=diff3). **J)** Lipidomics performed on enriched mitochondrial fractions from hDANs where the percent lipid content is normalised to the healthy control for cholesterols (left panel), cholesteryl ester (middle panel) and cardiolipin (right panel) (n=diff 3). **K)** Lipidomics performed on whole hDANs where the percent lipid content is shown for those lipid species downregulated in both PINK1 KO clones (left panel) and those lipid species upregulated in both PINK1 KO clones (right panel) compared to control. Two way ANNOVA comparing genotype and lipid.

The mitochondrial membrane potential (ΔΨm) of PINK1 KO hDANs is reduced by approximately 15% in the basal state and is less responsive to mitochondrial toxins in live cell imaging (Figure 2D). The phenotype was confirmed using live cell flow cytometry methods (Figure S1D). PINK1 KO hDANs do not show any significant oxidative stress, with regards to mitochondrial reactive oxygen species (ROS) (Figure 2E). Additionally, cytosolic ROS (Figure S1E), lipid peroxidation (Figure S1F), antioxidant status and glutathione depletion (Figure S1G) are also unaffected by PINK1 KO.

A metabolic shift away from oxidative phosphorylation is accompanied by ∼50% inhibition of the iron-sulfur cluster, TCA enzyme Aconitase (Figure 2F), which is in line with previous work *in vivo*^6^ and suggestive of upregulated citrate synthase and increased shuttling of citrate out of the mitochondria. Downstream in the TCA cycle, α-ketoglutarate dehydrogenase enzyme activity (αKGDH) is unaffected by PINK1 KO (Figure S1H). Significantly increased NAD^+^ (Figure 2G) but not NADH (Figure S1I) reflects metabolic shifting as an adaptive response.

We performed RNA sequencing in PINK1 KO hDANs in the basal state or following depolarization of the mitochondrial membrane with the ionophore Valinomycin to identify genes and pathways relevant to PINK1. The heatmap illustrates the most differentially regulated genes (log2 fold change) in both PINK1 KO hDAN clonal lines compared to their healthy control (Figure S2A). Pathway analysis comparing genotype revealed enrichment for genes involved in lipid synthesis and storage such as ELOV2/6 (long chain fatty acid (FA) synthesis), CYP27B1/CH25H (cholesterol regulation), FABP7 (FA binding and metabolism), COL1A2/COL4A4/COL4A2 (adipogenesis and fat storage), VCAM/LAMA1 (adhesion molecules involved in lipid metabolism and immune response) (Figure 2H). Interestingly, ionophore treatment of control neurons induces predominant expression changes in genes involved in amino acid metabolism, gluconeogenesis, glycolysis, TCA and iron homeostasis (Figure 2H).

Following on from these findings, we questioned whether lipid synthesis might be required for regeneration and biogenesis of mitochondrial membranes in the absence of PINK1 and asked whether specific lipids might be affected. We first measured cholesterol levels in whole cell preparations (Figure 2I) and then performed lipidomics on crude mitochondrial preparations (Figure 2J) and whole hDANs (Figure 2K). Cholesterol levels are increased by approximately 30% in PINK1 KO hDANs (Figure 2I), accompanied by a small increase in mitochondrial enriched samples which is not statistically significant (Figure S2B). Cholesterol and cardiolipins (CL) (Figure 2J) are also elevated in the crude mitochondrial preparations from PINK1 KO hDANs but the difference again is not significant. Lipidomics performed in whole hDANs showed both up and down regulation of lipid species in PINK1 KO lines (Figure S2C). Two way ANNOVA analysis of certain phosphatidylethanolamines (PE), phosphatidylcholines (PC), sphingomyelins (SM), CL and a ceramide b species where PINK1 KO hDANs have less than control, revealed significance but only for one of the two PINK1 KO clonal lines (Figure 2K, left panel). Upregulated lipid groups were significantly different to controls in both PINK1 KO clones. These lipids include: phosphatidyinositols (PI), PE, PS, PC, SM, CL and Bis(monoacylglycero)phosphates (BMP) (Figure 2K, right panel). Increased levels of phosphatidylethanolamines were significantly upregulated in statistical analysis of the individual lipid groups from whole hDAN analyses. Taken together, these data suggest a significant shift towards lipid abundance, which does not significantly alter the mitochondrial lipid pool. Cholesterol is most abundant at the plasma membrane and mitochondria only account for a small proportion of total cell cholesterol. Insulin-induced gene 1 protein (INSIG) and super conserved receptor expressed in brain (SREB) regulation at the ER may be relevant but requires much further work (especially the regulation of INSIG). Nevertheless, we could not detect any obvious differences in SREB co-localization with nuclei (Figure S2D). Total cholesterol levels were significantly increased in the striatum of four month old PINK1 KO mice (Figure S3C).

### PINK1 mechanisms switch during neuronal differentiation involving the phosphorylation of substrates at mitochondria and endoplasmic reticulum

We hypothesized that PINK1 could have additional or overlapping roles in mitochondrial quality control acting via substrates at the mitochondrial outer membrane (MOM), Endoplasmic reticulum (ER) or mitochondrial associated membranes (MAM) that could explain the metabolic changes observed. First we measured the distances between mitochondria and ER using quantitative BRET signals and found that PINK1 KO significantly tightens the connections in NPCs which loosen in post-mitotic mature hDANs compared to controls (Figure 3A).

**Figure 3.**
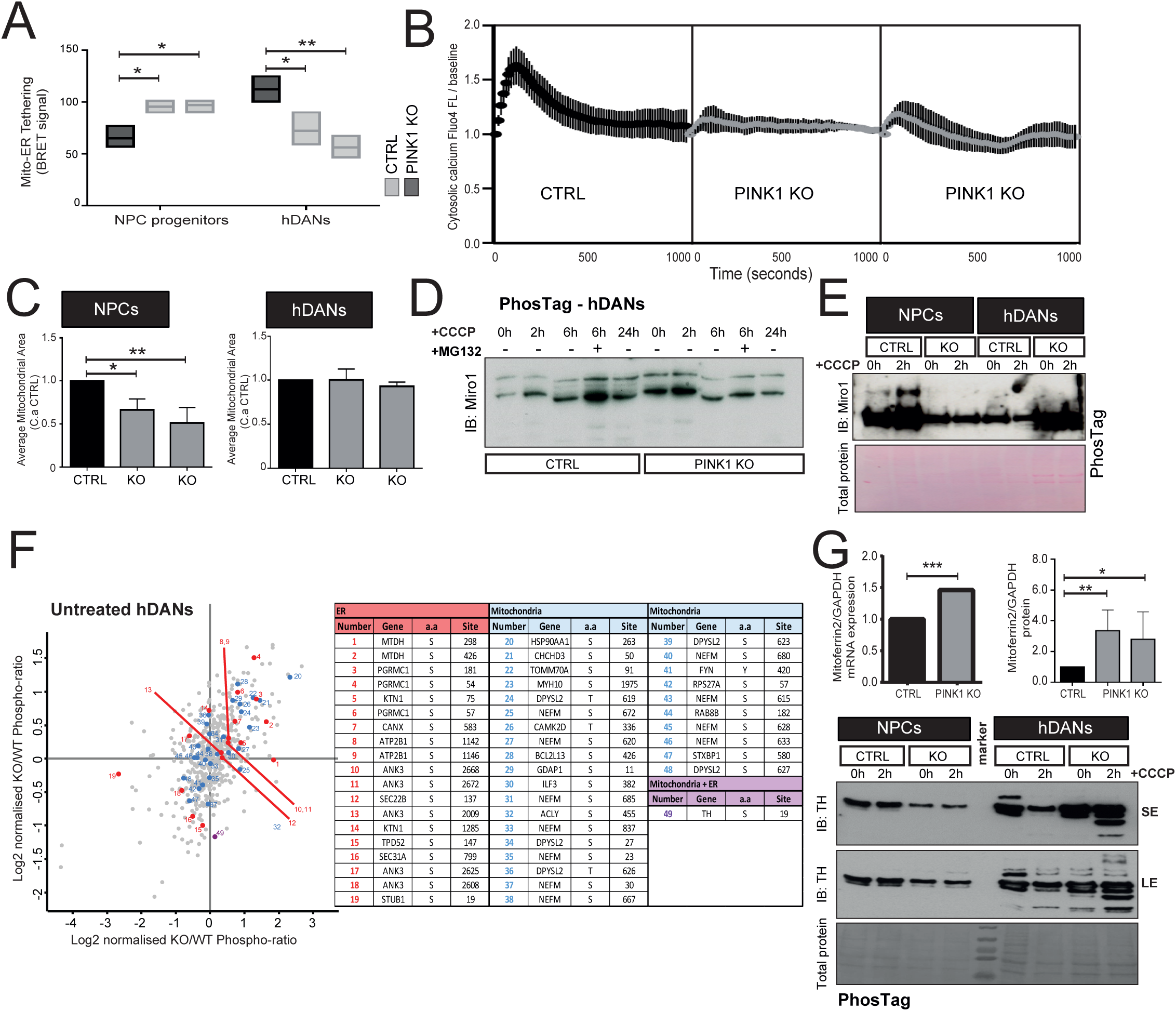
**A)** Mitochondrial-ER tethering (Mito-ER tethering) in NPC neuroprogenitors and hDANs measured by BRET signal which is increased as mitochondria and endoplasmic reticulum (ER) are closer in proximity (n=3 NPC independent experiments and hDAN differentiations). **B)** Cytosolic calcium levels in response to the addition of thapsigargin during live cell imaging measured by quantifying Fluo4 dye fluorescence in hDANs. The total corrected cell fluorescence (accounting for cell area) was normalised in each image to the baseline measurement before addition of thapsigargin after incubation with EGTA (n=diff4). **C)** Increased mitoferrin 2 gene expression (left panel) and protein abundance (right panel) normalised to GAPDH (n=diff3). **D)** Time course of Miro1 phosphorylation status following 10µM CCCP to stabilise PINK1 and MG132 to block Miro1 degradation using PhosTag SDS-PAGE (representative blots, n=diff3). **E)** Phosphorylation status of Miro1 in NPC neuroprogenitors and hDANs where CCCP is used to depolarise the MOM and stabilise PINK1 (representative blots, n=diff3). **F)** Plot of the most differentially phosphorylated proteins in enriched mitochondrial fractions from hDANs untreated or treated with Simvastatin. Log2 ratio of phosphor-proteins is normalised to the total proteome in each case (n=diff1). **G)** Mitoferrin 2 gene expression and protein levels relative to GAPDH (n=diff3) (upper panel). Phosphorylation status of tyrosine hydroxylase (TH) in NPC neuroprogenitors and hDANs where CCCP is used to depolarise the MOM and stabilise PINK1 (representative blots, n=diff3).

Broader distance between mitochondria and ER reflects the glycolytic, lipid rich metabolic state observed in caloric excess and PINK1 KO hDANs. Furthermore, the ER of PINK1 KO hDANs fail to release normal amounts of calcium in response to thapsigargin treatment (Figure 3B). Tighter mitochondrial contacts in PINK1 KO NPCs is accompanied by significantly smaller mitochondria, suggestive of fragmentation (Figure 3C). Interestingly, PINK1 KO hDANs show no mitochondrial fragmentation (Figure 3C) nor any other changes in mitochondrial morphology compared to controls using live cell microscopy (Figure S2E).

Next we studied the phosphorylation status of Miro1, a known substrate of PINK1 which is important for mitochondrial-ER contact, mitochondrial morphology and mitochondrial movement. Phosphorylation of Miro1 is largely undetectable unless an ionophore is used to initiate PINK1 accumulation at the MOM. In control hDANs, phosphorylation events on Miro1 accumulate during CCCP treatment. Several phosphorylated Miro1 bands are absent in PINK1 KO lysates run on PhosTag gels (Figure 3D). PINK1 KO affects the phosphorylation of Miro1 more strongly in NPCs than in hDANs (Figure 3E). Grp75 and TOM70 may also be phosphorylated in the presence of PINK1 and this may again be more important in NPC intermediates (Figure S2F).

To explain PINK1 KO phenotypes in hDANs, we looked for putative PINK1 targets using quantitative phospho-proteomics on crude mitochondrial preparations. We normalized phospho-peptide abundance to the non-phospho proteome in the same samples and analyzed untreated hDANs and those treated with the cholesterol lowering drug Simvastatin to identify any putative substrates relevant to PINK1 regulation of lipid metabolism. We ranked phospho-protein hits based on those with the greatest log2 fold change and removed those that did not replicate in both PINK1 KO clones (Figure S2G). To focus on interactions that may occur at the mitochondrial contact sites, we then listed phospho-proteins by the GO terms; mitochondria, ER, mitochondria and ER (Figure 3F). We identified GAP43 (Neuromodulin) which is a protein associated with calcium signaling and neuronal development and validated biochemically by PhosTag gels (Figure S2H). Of note, proteins related to iron and hormone regulation such as PGRMC1, an ER protein which is part of the ferrechelotase complex, was also validated (Figure S2H). Mitoferrin2, a protein involved in iron import into mitochondria is upregulated in PINK1 KO hDANs (Figure 3G). We monitored the mitochondrial uptake of labelled proteins and found no significant impairment in the absence of PINK1 (Figure S3D).

Finally, we identified that PINK1 could be important for the phosphorylation of tyrosine hydroxylase (TH) at serine 19 (Figure 3F), a protein filtered from phospho-proteomics data based on its GO terms (it localizes to both mitochondria and ER). We biochemically validated the event in PINK1 KO NPCs and hDANs. Missing and altered phospho-TH banding is observed in PINK1 KO hDANs on PhosTag gels (Figure 3G). PINK1-dependent phosphorylation of TH occurs in untreated hDANs and depolarization of the mitochondrial membrane with CCCP does not increase phosphorylation at the same band (Figure 3G), suggesting that PINK1 may not directly phosphorylate TH and membrane depolarization may be more important.

### PINK1 KO induced membrane rigidity alters the distribution and regulation of the dopamine transporter

Upregulation of Rab3B expression was detected following RNA sequencing by qRT-PCR (Figure S3A). Rab GTPases are known PINK1 targets ^72^ and Rab8A is involved in recycling cholesterol to the plasma membrane^73^. We therefore questioned whether deregulation of cholesterol reported here could impact lipid membranes crucial for synaptic function in neurons. We used dynamic light scattering to measure the size and heterogeneity of membrane particles following mechanical homogenization of hDANs in the absence of detergents. We found PINK1 KO hDANs contain more disperse, larger particles compared to their healthy control (Figure 4A). Next we fractionated hDAN lysates by density and used marker proteins to identify the soluble fractions (8-14) and the cholesterol-rich floating fractions (3-7) marked by flotillin (Figure 4B). We assessed the distribution of several membrane-bound and cytosolic proteins in precipitated fractions from hDANs. Quantification of protein levels per fraction revealed that the lipid raft marker flotillin is distributed more heavily in the floating fractions indicative of increased cholesterol rich rafts in PINK1 KO hDANs (Figure 4C, upper panel). The distribution of ERK1/2, which is an abundant cytosolic protein is unaffected by PINK1 KO (Figure 4C, bottom panel) suggesting that membrane bound proteins may be particularly affected by changes in cholesterol. Since the dopamine transporter DAT is known to be negatively regulated by its phosphorylation in cholesterol rich lipid membrane rafts^74^ reviewed in^75^, we then assessed the distribution of phosphorylated DAT and total DAT. PINK1 KO leads to more phosphorylated DAT in less soluble fractions (Figure 4D), which leads to increased DAT internalization. We treated hDANs with 1mM β-cyclodextrin to deplete cholesterol, break up lipid rafts and push proteins into the soluble fractions as a technical control (Figure 4E). In collaboration with the authors of Jazaro et al. (submitted to NCB together with this manuscript) who had independently found that sub-lethal doses β-cyclodextrin reverse PINK1-specific phenotypes in patient derived models via unbiased compound screening, we then used nanomolar concentrations of β-cyclodextrin as a chronic treatment in the maturation media of hDANs and found that it was capable of reversing the reduced mitochondrial membrane potential previously observed (Figure 4F) also reduced utilization of exogenous fatty acids as an energy source (Figure 4G).

**Figure 4.**
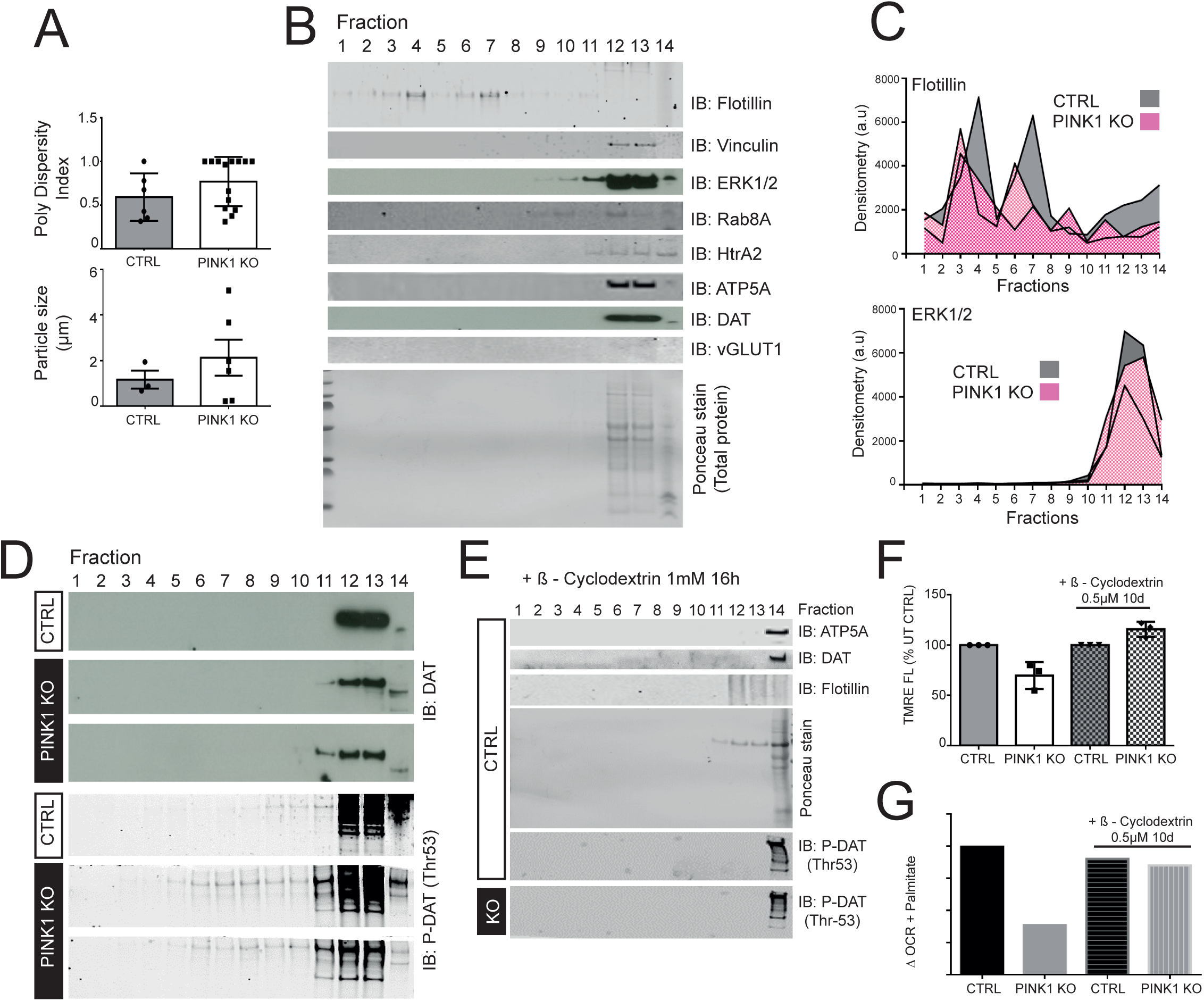
**A)** Properties of particles in hDAN homogenates; dispersity (heterogeneity of shape and size) index measured by dynamic light scattering (upper panel) where a poly dispersity index (Pdi) of 1.0 reflects a homogenate that contains extremely heterogeneously sized particles not considered suitable for particle size measurement. Particle size was calculated for those homogenates with a Pdi of 0.75 or under using dynamic light scattering (n=diff3). **B)** Sucrose gradient fractionation of total hDAN lysates followed by protein precipitation and SDS PAGE to detect markers of the soluble and lipid raft floating fractions (representative blots, n=diff3). **C)** Graph showing the amount of quantified flotillin (lipid raft marker) and ERK1/2 (cytosolic marker) distribution across the 14 fractions. **D)** Distribution of dopamine transporter (DAT) and Phospho-DAT (P-DAT Thr53) throughout the hDAN fractions (n=diff3). **E)** Acute 1mM ß-cyclodextrin treatment depletes cholesterol and solubilises marker proteins in hDANs (n=diff3). **F)** Chronic 500nM ß-cyclodextrin treatment of hDANs during neuronal maturation restores mitochondrial membrane potential measured by TMRE fluorescence using flow cytometry (n=diff6). **G)** Chronic 500nM ß-cyclodextrin treatment of hDANs during neuronal maturation restores oxygen consumption (respiratory) response to utilise exogenous fatty acids as an energy source.

### PINK1 knockout significantly affects dopamine uptake and neurotransmission

We observed differences in DAT distribution and TH phosphorylation in PINK1 KO hDANs. We also observed altered expression of genes involved in dopamine metabolism and synaptic function. Of particular interest, upregulation of pyridoxine-5’-phosphate oxidase (PNPO), which is important for the synthesis of amino acids and dopamine via AADC, and of TH, which is important in dopamine synthesis, where observed (Figure S3B and Figure 1C). We therefore measured amine neurotransmitter metabolites in control and PINK1 KO hDANS. L-DOPA is routinely used in cell culture models to feed the system a substrate in order to measure dopamine (DA) and DA flux accurately using HPLC. The PINK1 KO hDANs have significantly reduced DOPAC and DA levels (Figure 5A). The ratio of DOPAC/DA was also reduced but not HVA/DA suggesting the importance of the pre-synapse (Figure 5B). We controlled for amount of dopaminergic neurons in the cell cultures by monitoring TH on mRNA and protein level in each experiment, which was not decreased in PINK1 KO hDANs (Figure 5C). DA levels inside hDANs and in the media could not be replenished by blocking degradation (Figure 5D). MAO A and B enzyme activity and MAO-A protein levels were not significantly affected by PINK1 KO (Figure 5E). However, since MAO levels are normally highly fluctuating, a possible functional role of the enzymes need to be further addressed.

**Figure 5.**
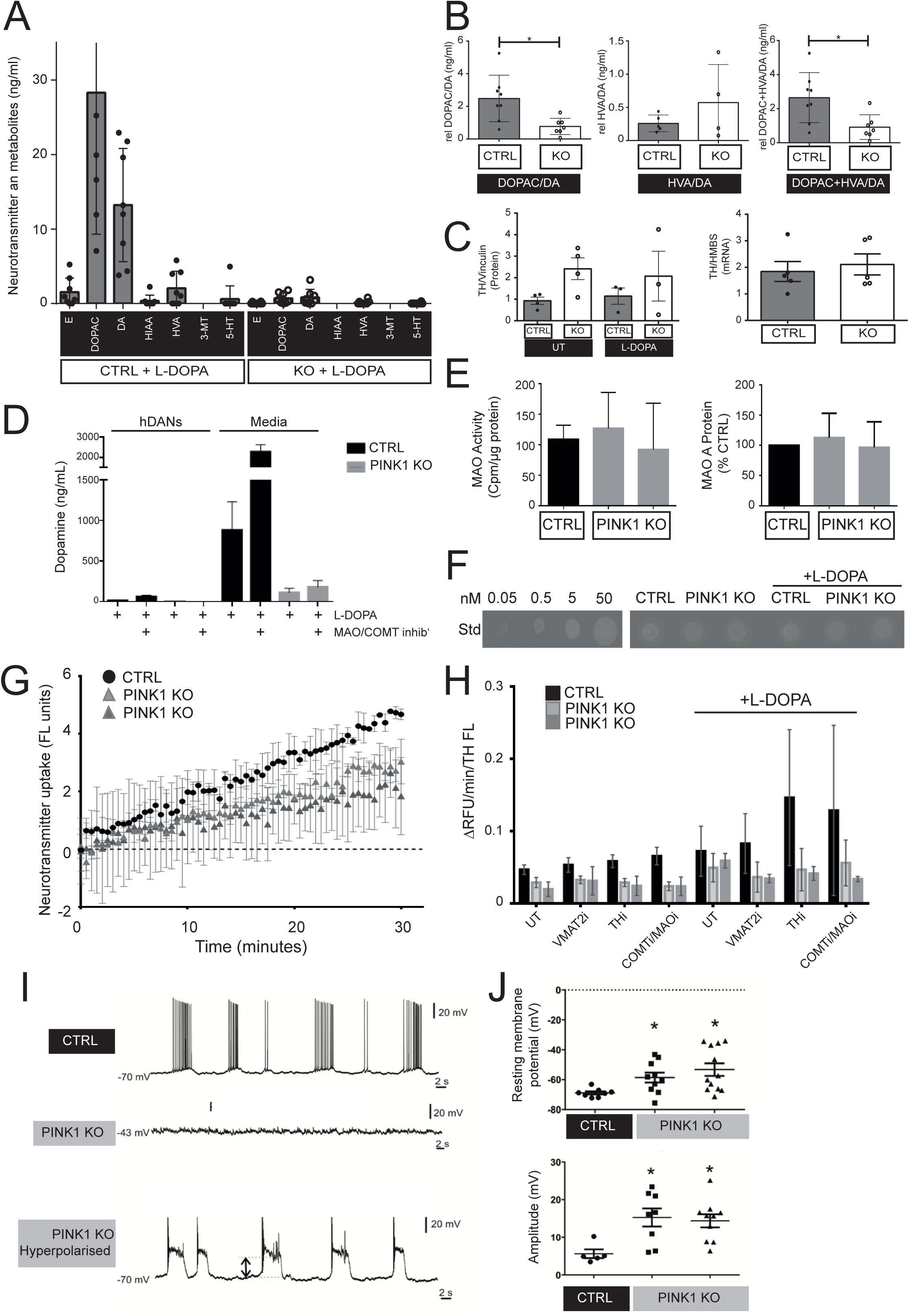
**A)** Concentration of neurotransmitters and metabolites epinephrine (E), 3,4-Dihydroxyphenylacetic acid (DOPAC), Dopamine (DA), 5-Hydroxyindoleacetic acid (HIAA), Homovanillic acid (HVA), 3-Methoxytyramine (3-MT) and 3-(2-aminoethyl)-1 H -indol-5-ol /Serotonin (5-HT) in hDANs treated with L-DOPA for 24h (n=diff5). **B)** Ratios of DOPAC/DA, HVA/DA and DOPAC plus HVA/DA. **C)** Tyrosine hydroxylase (TH) protein levels (left panel) and gene expression (right panel) in hDAN aliquots from each experiment/hDAN differentiation shown in A) and B). **D)** Dopamine concentration in hDANs and in cell culture supernatants treated with L-DOPA for 24h with or without inhibition of dopamine degradation via COMT and MAOA/B (MOA/COMT inhib’) n=diff3). **E)** MAO A/B activity in hDANs measured by the deamination of radiolabelled tyramine (left panel) and MAO-A protein levels normalised to GAPDH marker and the healthy control (right panel) (n=diff3). **F)** Catecholamine oxidation in hDANs with or without L-DOPA treatment for 24h (representative blots, n=diff3). **G)** Neurotransmitter uptake in hDANs measured by the fluorescence of the labelled amine converted inside hDANs only (n=diff3). **G)** Quantification of neurotransmitter uptake in hDANs with or without acute inhibition of TH activity (THi), inhibition of VMAT2 activity (VMAT2i), inhibition of COMT and MAOA/B activity (COMTi/MAOi) with or without 24h L-DOPA treatment. The rate of uptake is normalised to the amount of TH staining in the well to account for hDAN number (n=diff3). **I)** Trace activity of (TH positive) hDANs patched and unstimulated (two upper panels) and PINK1 KO hDANs following hyperpolarisation (n=diff3). **J)** Resting membrane potential (upper panel) and amplitude (lower panel) of hDANs (n=diff3).

Neurotransmitter uptake was significantly impaired in PINK1 KO hDANs (Figure 5F and Figure 5G) and this could not be rescued by inhibition of uptake in the vesicles (VMAT), DA synthesis via TH or DA degradation (COMT/MAO) (Figure 5H). We measured the amount of oxidized catecholamines and did not observe any differences between PINK1 KO hDANs and their control with or without L-DOPA treatment (Figure 5H). Therefore, DA loss cannot be explained by oxidation but conversion to neuromelanin is still possible but difficult to detect in iPSC-derived 2D hDAN cultures.

Next we tested the electrophysiological properties of PINK1 KO hDANs using patch clamp recordings. Healthy hDANs showed consistent burst firing patterns, while PINK1 KO hDANs showed 50% spontaneous bursting or 50% no spontaneous action potentials (Figure 5I). PINK1 KO hDANs have a significantly elevated mean average resting membrane potential above −50mV (Figure 5J). PINK1 KO hDANs generated action potentials when we artificially lowered their membrane potential to −70 mV, the normal resting potential of healthy cells. In this state PINK1 KO hDANs were able to produce spontaneous bursting behavior similar to the healthy controls. However, these PINK1 KO hDANs have an increased amplitude of the bursting (5.64 ± 1.05 control vs 15.26 ± 12.26 PINK1 KO and 14.39 ± 1.63 PINK1 KO) and often showed depolarization blocks within the bursting (Figure 5I, lower panel). PINK1 KO hDANs have an abnormal and hyper-excitable phenotype. Jazaro et al. (submitted together with this manuscript) found that abnormal firing patterns in PINK1 patient derived models can be rescued by low doses of β-cyclodextrin, suggesting that plasma membrane fluidity may be an important factor in PINK1 PD etiology.

## Discussion

It is widely accepted that wild type PINK1 acts as a mitochondrial sensor, an initiator of stress responses and is important for mitochondrial quality control. We show here that PINK1 is not essential for the maintenance of mitochondrial networks or mitochondrial respiratory function in mature human neurons, which is in line with prior evidence that mitophagy can occur independently of PINK1 ^37, 56, 76^.PINK1 orchestrates a network of fine tuning mechanisms such as PINK1-Parkin mediated mitophagy, inflammation and mitochondrial movement^21, 23, 66, 77^. Here we show that PINK1 is also important for mitochondrial metabolism. Metabolic control within the mitochondrial matrix might be imposed by PINK1 via several mechanisms; firstly, via inner mitochondrial targets such as Aconitase, CI or TRAP1 which have all been previously proposed^6, 70, 78^. PINK1 could indirectly regulate the mitochondrial import of iron or other molecules required as enzymatic co-factors. This is an interesting topic that requires additional work to go beyond protein import into mitochondria. PINK1 is known to regulate Miro1, which is a MOM protein crucial for calcium sensing, movement and mitophagy. It would be interesting to detangle the boundaries between mitophagy and mitochondrial-ER contact regulation since many PINK1-Parkin targets have functions spanning both processes.

Mitochondrial fragmentation and compensation occurring in developing neurons, which is overcome for the survival of the mature neurons supports the model that fission protects healthy mitochondrial domains from elimination when unchecked by the PINK1-Parkin pathway^79^, PINK1-independent mitophagy and alternative mitochondrial quality control pathways such as piecemeal mitophagy^80^ and mitochondria derived vesicles^81^. Here, the synthesis of lipids for renewal of the membranous mitochondrial network may be relevant.

We propose that neurons lacking PINK1 perform a citrate shunt catalyzing Acetyl CoA production and fatty acid synthesis. Mitochondrial-ER contacts in developing neurons could be favored to increase mitochondrial capacity by via ß-oxidation^82^. Here oxidative damage is absent and the increased NAD+ pool further underscores the relevance of an adaptive response in early differentiation.

In mature neurons distant mitochondrial-ER contacts reflect a high fed state and hyperlipidemia ^82^. In an extensive metabolomics study of PD patients, changes in long chain acyl carnitines and their role in β-oxidation were highlighted as a robust PD biomarker^83^ and lipid regulating proteins were the most significant modifiers of PINK-mediated mitophagy in another unbiased screen^59^.

Cholesterol is one of the major components of lipid rafts which are crucial at the synapse and post synapse. Lipid rafts control the movement and modification of receptors and transporters but are also involved in endocytosis, exocytosis and vesicular transport. Previously Rab8A has been identified as a target of PINK1^72^ which is involved in membrane transport but also in cholesterol recycling^73^. This pathway may be crucial for PINK1-specific mechanisms outside of mitochondria and the link to the synapse.

Altered dopamine uptake and calcium mismanagement contribute to the abnormal firing patterns and very high resting membrane potentials in PINK1 KO hDANs. We cannot rule out that low dopamine metabolites are caused by the additional impact of catechol estrogens and other lipid hormones on neurotransmitter synthesis ^84, 85^ or dopamine synthesis via AADC which requires PLP, a product of the vitamin B6 salvage pathway which is significantly down regulated in PINK1 KO hDANs at the enzyme PNPO. PLP is abundant in the base medias used for neuronal differentiations and therefore a different approach is required. PLP is crucial for amino acid, nucleotide and neurotransmitter synthesis^86^. Enhancing nucleotide metabolism was previously found to protect against PINK1-PD^87^ and therefore could be an entry point for therapeutics. PINK1 specific modification of TH at the mitochondrial membrane or at MAMs provides addition burden to dopaminergic neurons and the PD gene product α-synuclein has previously been implicated in TH regulation at mitochondrial-ER contact sites^88^. Parkin is known to suppress MAO^89^, a mitochondrial anchored enzyme responsible for dopamine degradation and mitochondrial DNA damage^90^ and which is implicated in the pathogenesis of PD^91^. Our data underscores the relevance of dopamine regulation at the mitochondria in human neurons.

Here, the solubilizing agent ß-cyclodextrin (BCD) depleted cholesterol and therefore the lipid rafts (according to^92^) and was protective at low doses. Jarazo and colleagues (in the accompanying paper in this submission) found that BCD rescues mitochondrial and neuronal phenotypes in PINK1 patient-derived models. The mechanism of BCD action could be relevant for other forms of PD or neurodegeneration. Indeed, mild membrane cholesterol reduction also impacts the cleavage of APP mediated by APP trafficking and partitioning into lipid rafts^93^. Although Simvastatin use for brain related diseases is complicated by the dissociation of cholesterol metabolism in the brain (reviewed in^94^), data taken from the Norwegian medical database shows Simvastatin is associated with decreased risk of Parkinson’s disease^95^. Further mechanistic work will determine whether BCD could be beneficial in other human neuronal models of Parkinson’s disease.

## Materials and Methods

### Ethics statement

Human samples were obtained with consent and prior ethical approval at The University of Tübingen and the Hertie Institute for Clinical Brain Research Biobank.

Animal experiments were carried out in strict accordance with the recommendations in the Guide for the Care and Use of Laboratory Animals of the European Union and of the Federal Republic of Germany (TierSchG). The protocol was approved by the Institutional Animal Care and Use Committee (Ausschuss für Tierversuche und Versuchstierhaltung, ATV) of the Helmholtz Zentrum München. All efforts were made to minimize suffering.

### Generation of iPSC and PINK1 Knockout

Human iPSCs were cultured in self-made E8 media on Vitronectin (VTN-N, Gibco) coated cell culture dishes. iPSCs from a healthy individual that were previously characterized ^96^ were transfected with a TALEN and a homologous construct for *PINK1* exon1 using an Amaxa Nucleofector II with the Stem cell Nucleofection Kit (both from Lonza). The transfected iPSCs were plated on VTN-N –coated 10cm dishes in E8 medium containing 10 µM ROCK inhibitor Y27632. Homologous recombined iPSC colonies were selected with 250 µg/ml G418 (Biochrom) or 10 µgml-1 Blasticidin (InvivoGen) in the second round of TALEN transfection and re-plated in 12-well plates. Resistant iPSC colonies were characterized by sequencing, qRT-PCR and Western blot to confirm successful homozygous gene knockout. TALENs were designed with the online tool TAL Effector Nucleotide Targeter 2.0 (Cornell University) and generated using a cloning protocol adapted from ^97^. The following RVD sequences were used for the TALEN monomers: HD HD NI NH NH NG NH NI NH HD NH NH NH NH HD and NI NH HD NG HD HD NH NG HD HD NG HD HD NH HD. Colony-PCR after the first TALEN reaction was conducted with the primers pCR8_F1 (5’-TTGATGCCTGGCAGTTCCCT-3’) and pCR8_R1 (5’-CGAACCGAACAGGCTTATGT-3’). After the second Golden Gate reaction the colony-PCR was performed with the primers TAL_F1 (5’-TTGGCGTCGGCAAACAGTGG-3’) and TAL_R2 (5’-GGCGACGAGGTGGTCGTTGG-3’).

### iPSC Differentiation into mature, human, mid-brain specific dopaminergic neurons (hDANs)

hDANs were generated from iPSCs via neuropregenitor (NPC) intermediates using a protocol adapted from (Reinhardt, Glatza et al., 2013b). Briefly, IPSCs were maintained in E8 medium. For the generation of embryoid bodies (EBs), iPSCs were cultured in ‘50:50 base medium’ (one to one mixture of DMEM Hams F12 (#FG4815 Biochrome/Millipore): Neurobasal® medium (#21103-049 Gibco/Thermo Scientific), 1X Penicillin/Streptomycin (Biochrome/Millipore), 1X GlutaMAX supplement Thermo Scientific), 1X B27 supplement (Gibco/Thermo Scientific) and 1X N2 supplement (Gibco/Thermo)) plus the addition of 10µM SB431542 (Sigma, SB), 1µM dorsomorphin, 3µM CHIR99021 (CHIR, Axon) and 0.5µM pumorphamine (PMA, Alexis) on uncoated 6-well cell culture plates. EBs were then transferred to Matrigel (Corning)-coated 6-well plates in NPC maintenance media (’50:50 base medium’ plus the addition of 150µM Ascorbic Acid (AA, Sigma), 3µM CHIR, 0.5µM PMA). After several passages NPCs were cultivated in NPC priming medium (‘50:50 base medium’ plus the addition of 150µM AA and 3µM CHIR99021). Differentiation of confluent NPCs was initiated by cultivation in a patterning medium for seven days (‘50:50 base medium’ plus the addition of 10ng/mL FGF8 (Peprotech), 1µM PMA, 200µM AA, 20ng/mL BDNF (Peprotech). The differentiating neurons were matured in maturation media (’50:50 base medium’ plus the addition of 10ng/mL BDNF, 10ng/mL GDNF (Peprotech), 1ng/mL TGFß-III (Peprotech), 200µM AA, 500 µM dbcAMP (Applichem) and 10µM DAPT (Sigma). Prior to all mitochondrial function experiments (ATP assay, measurement of reactive oxygen species, MitoTimer, respiratory measurements, complex I activity assay, glutathione assay, lipid peroxidation, apoptosis, LDH release and mitochondrial membrane potential), maturation medium was replaced 24h before the experiment with ‘N2 medium’ (using 50:50 base media without B27 supplement or ascorbic acid) to reduce excessive amounts of antioxidants in the media that could quench oxidative stress or respiratory phenotypes.

### Animal work

Pink1-knock-out mice (Glasl et al., 2012) with 4–16 months of age were used and bred in accordance with the regulations from the government of Upper Bavaria. The mice are on kept on a C57BL/6J background. All mice were kept on an inverse 12-h light/12-h dark cycle (lights off at 18:00). Mice were provided with ad libitum access to standard chow and water. Mice were killed by cervical dislocation and the brain immediately removed. Thereafter single brain regions were dissected and the tissue shock frozen in liquid nitrogen.

### Quantitative Reverse Transcription PCR (qRT-PCR)

RNA was isolated from neurons using a RNeasy Mini Kit (QIAGEN), including the on-column DNA digestion step. A one-step qRT-PCR was performed on 0.1-1µg RNA (equalized to the same imput amount) with the QuantiTect SYBR Green RT-PCR Kit (QIAGEN) on a LightCycler®480 (Roche). The relative expression levels were calculated with the 2^−Δ^ method, based on a biological reference and housekeeping genes (GAPDH and HMBS) for normalization. The following primer sequences were used

### Immunofluorescence staining

hDANs were cultivated on Matrigel-coated glass coverslips and fixed with 4% (w/v) paraformaldehyde (PFA, Sigma) in phosphate buffered saline (PBS) for 15 min at room temperature (RT). Permeabilisation with ice-cold, neat methanol for 5 min at - 20°C. After washing with 0.01% (v/v) Tween in PBS (PBS-T), the fixed hDANs were blocked with 5% (v/v) normal goat serum in PBS-T for 1h at RT. Afterwards the cells were washed again and incubated overnight at 4°C with the primary antibody in 2.5% (v/v) serum in PBS-T. On the following day the cells were incubated with the secondary fluorescent antibody (#A32721, #A11070, #A21449, #A11010 all from Molecular probes/Thermo Scientific) for 2h at RT in darkness and nuclei stained with DAPI (Sigma). Coverslips were mounted in mounting medium (Dako) on glass slides. Immunofluorescence was imaged using an AxioVert fluorescence microscope (Zeiss). MAP2 (AbCam ab5392) (n=diff5), TH (AbCam ab112) (n=diff5), FOXA2 (Millipore AB4125) (ndiff 3), DAT (Millipore MAB369) (nDiff3).

### LDH Assay

LDH release was measured using the Pierce LDH Cytotoxicity Assay kit (Thermo Scientific). hDANs were seeded on a Matrigel coated 96 well plate. hDANs were treated in triplicates with and without 50 nM and 100 nM rotenone in N2 medium for 24h at 37 °C with 5 % CO_2_. 110µL of the celcu l lture medium was transferred to an empty 96-well plate to obtain a spontaneous release fraction. Lysis buffer was added to the cell plate and then both 96-well plates were incubated for 45 minutes at 37 °C with 5 % CO_2_. Both plates were centrifuged for 5 min at 3220 rpm at RT. 30µL sample from each cell line, each condition and each technical replicate was transferred to a 96-well ELISA reader plate (Cellstar) and the reaction started, measured and stopped according to the manufacturer’s recommendations. LDH activity was calculated by subtraction of the 680nm absorbance from the 490nm absorbance values. Cytotoxicity was determined by dividing the LDH activity from the spontaneous release fractions by total LDH activity from the lysed samples. Increased LDH release caused by cell damage and apoptosis.

### SDS-PAGE and Western blotting

All cell lysates were prepared in RIPA buffer (Sodium chloride 150mM, Tris-HCL 50mM, Sodium dodecyl sulfate 0.10% (w/v), Sodium deoxycholate 0.50% (w/v), Triton-X-100 1% (v/v)) containing 1X concentration of phosphatase inhibitor cocktail (Complete, Roche) and phosphatase inhibitor cocktail (PhosSTOP, Roche). Briefly, the lysis buffer was added directly to washed cells in dishes or washed cell pellets and kept at 4 °C. Needles were used to further homogenize the lysates (9 passes 20G, 9 passes 27G) and incubated 30 minutes on ice. Insoluble nuclear material was removed after centrifugation at 14,000 rpm for 10 minutes. Proteins were electrophoresed on self-made acrylamide gels or pre-cast Bis-Tris gels (Thermo Scientific) and transferred to nitrocellulose membranes using the iBlot device (Thermo Scientific), with the exception of very large target proteins or heavily lipidated proteins, in which case wet blotting with PVDF membranes were used. Total protein stain Ponceau (Applichem) was used to assess transfer and loading and the PageRuler plus pre-stained protein ladder (Thermo Scientific) for kDa range. Antibodies against mAconitase (BD Biosciences), β-Actin (Sigma Aldrich), GAPDH (Invitrogen, Thermo Scientific) Tom20 (Santa Cruz Biotechnology), LC3I/II (Novus), PGC1α (AbCam), Miro1 (Sigma Aldrich), OPA1 (Novus Biologicals), TH (Millipore), Flotilin (Santa Cruz), Vinculin (Sigma-Aldrich), Rab8A (Cell Signaling Technology), HtrA2 (R&D Systems), ATP5A (AbCam), DAT (Millipore), P-Thr53 DAT (Thermo Fisher), vGLUT1 (Stressmarq), MFRN2 (Thermo Fisher), GRP75 (Santa Cruz), GAP43 (Novus) and PGRMC1 (Proteintech), ERK1/2 and P-ERK1/2 (Cell Signaling Technolgy) and mitobiogenesis antibody (containing SDH, GAPDH and COX, AbCam) were used. Secondary antibodies were purchased from GE Healthcare (HRP-conjugated) and from LiCOR (α-rabbit and mouse Alexa Fluophor™680, α-rabbit and mouse Alexa Fluophor™800). Fluorescence detection and analysis were performed using a LI-CORE blot scanner and Image Studio™ Lite software. Densitometry from Western blot was performed using the Image J 1.41o software (Wayne Rasband; National Institutes of Health, USA).

### Autophagy and mitophagy induction

Autophagic flux was induced in hDANs using 50 mM ammonium chloride for 1h, 1µM Valinomycin for 24h. Mitophagy was induced in hDANs using; 10µM CCCP for 2h, 4h, 6h, (6h+ 10µM MG132) and 24h or 1µM Valinomycin for 24h.

### Mitochondrial turnover using MitoTimer^™^

Mitotimer is a fluorescent reporter suitable for the investigation of mitochondrial turnover. Based on oxidation state, the fluorescent protein (dsred mutant) targeted to the mitochondria shifts its emission spectra from green to red as the protein matures.^98^. Approximately 100,000 hDANs were seeded per 24-well well on coated coverslips in maturation medium.

For the analysis the IMARIS software was used. hDANs were transfected with pMitoTimer a gift from Zhen Yan (Addgene plasmid # 52659; http://n2t.net/addgene:52659; RRID:Addgene_52659) using Fugene HD (Promega) according to manufacturer’s instructions. The cells were fixed 8, 24 and 48 h post-transfection with 4% (v/v) PFA in PBS for 10 min RT. Transfected neurons were imaged for both green (Ex/Em, 488nm/518nm) and red channels (Ex/Em, 543nm/572nm) using a LSM 510 confocal (Zeiss). Z-Stack pictures were analyzed using Imaris 8.3.1 (Bitplane). The interpretation of the red:green MitoTimer ratio as an indication of mitochondrial turnover was according to PMID:4128932.

### Respiratory analyses

For the basic mitochondrial stress test, OCR and ECAR were measured in NPCs and hDANs using a Seahorse™ XF96 Extracellular Flux Analyzer. 24 h. Cells were seeded in Matrigel coated Seahorse cell plates 24-48h prior to the experiment. During the experiemnet cells were treated sequentially with 1µM Oligomycin (Santa Cruz Biotechnology), 5µM FCCP (Santa Cruz Biotechnology) and 1µM Antimycin A (Santa Cruz Biotechnology)/ 1µM Rotenone (Sigma) in order to perform a mitochondrial stress test where the base medium contained glucose, pyruvate and glutamine. Normalisation was performed by counting the number of cells per well of the plate using a high content imager from BD Biosciences. Following the respiratory analysis, the media is removed and the cells are fixed with 4% (w/v) PFA containing Hoescht stain (1:10,000) for 5 minutes. Then washed and stored in PBS for imaging.

For the Fatty Acid Oxidation (FAO) test, OCR and ECAR were measured in hDANs seeded on Matrigel coated seahorse cell plates 24-48h prior to the experiment with 30,000-40,000 cells per well. Briefly, hDAN media was exchanged for FAO media (Seahorse base medium containing 2.5mM glucose, 5mM HEPES, 0.5mM Carnitine, pH7.4). After 20 minutes incubated in a non-CO2 incubator, BSA, or BSA-Palmitate was added to half of the plate in the presence or absence of Etomoxir (Eto) before starting the assay. Measurements were made in the basal sate before injection of oligomycin, followed by FCCP and finally rotenone plus antimycin A as in the mitochondrial stress test above.

### Flow cytometry experiments

hDANs were carefully washed in PBS and then treated with Accumax to remove them from the monolayer, quenched in PBS and centrifuged at 300 g for 5 minutes and then incubated in dye, buffer only or dye plus a control.

For early apoptosis; Annexin V-Pacific blue in Annexin V binding buffer (both from BioLegend) or Annexin V-Pacific blue plus Staurosporine was used.

For mitochondrial membrane potential, 200 nM Tetramethylrhodamine, Ethyl Ester, Perchlorate (TMRE, from Thermo Scientific) in PBS or TMRE plus Carbonyl cyanide-p-trifluoromethoxyphenylhydrazone (CCCP) 10 µM was used.

For mitochondrial reactive oxygen species, 2 µM MitoSox™ (Thermo Scientific) in PBS or MitoSox™ plus 10µM Rotenone was used.

Cells were measured using a MACSQuant® automated flow cytometer (Mitenyi Biotechnology) according to their mean average fluorescence signal. All mean average fluorescence values were divided by the background fluorescence in the same channel in the same unstained cells to account for auto-fluorescence.

### Live Cell Kinetic Measurement of Mitochondrial Membrane Potential

Cells were seeded in Ibidi® dishes and the media exchanged for HBSS containing 200 nM TMRE stain (Thermo Scientific) for 15 minutes at 37°C with CO_2_. The TMRE was removed and replaced with 360 µl Hanks buffer. The cells were imaged using a Zeiss Inverted Confocal microscope at Excitation HeNe1, 543 nm and Emission LP 560 nm and brightfield for 20 × 4s cycles. Followed by the addition of 360 µL (0.25 mg/ml Oligomycin), measured for 20 × 4s cycles, 180 µL (10 µM Rotenone), measured for 20 × 4s cycles and 100 µL (10 µM FCCP) and measured for 20-40 × 4s cycles. Using Image J, each transfected cell (detected using ZsGreen-TRAP1) in each frame was analysed for TMRE fluorescence intensity, mean fluorescence and total area. The corrected total cell fluorescence (CTCF) over time was calculated using the formula: CTCF = fluorescence intensity-(cell area × mean background fluorescence).

### RNA sequencing

Approximately 5 million hDANs were lysed in 350ml RTL buffer and homogenized using QIAshredder® homogenizer (Qiagen). RNA isolation was performed using RNeasy Mini Kit (Qiagen). RNA was eluted in 30μl RNase-free water. RNA quality was assessed with an Agilent 2100 Bioanalyzer and the Agilent RNA 6000 Nano kit (Agilent). Samples with very high RNA integrity number (RIN > 9) were selected for library construction. For polyA enrichment, a total of 200ng of total RNA was subjected to polyA enrichment and cDNA libraries were constructed using the resulting mRNA and the Illumina TruSeq Stranded mRNA kit (Illumina). Libraries were sequenced as single reads (65 bp read length) on a HighSeq 2500 (Illumina) with a depth of >22 million reads each.

Library preparation and sequencing procedures were performed by the same individual and a design aimed to minimize technical batch effects was chosen.

Read quality of raw RNA-seq data in FASTQ files was assessed using QoRTs (v1.2.37) to identify sequencing cycles with low average quality, adaptor contamination, or repetitive sequences from PCR amplification. Reads were aligned using STAR (v2.5.3a) allowing gapped alignments to account for splicing to the Ensembl Homo sapiens GRCh37 reference genome. Alignment quality was analyzed using ngs-bits (v0.1) and visually inspected with the Integrative Genome Viewer (v2.4.19). Normalized read counts for all genes were obtained using subread (v1.5.1) and edgeR (v3.24.3). Transcripts covered with less than 1 count-per-million in at least 5 out of 6 sample were excluded from the analysis leaving >13,000 genes for determining differential expression in each of the pair-wise comparisons between experimental groups.

#### Pathway Analysis

CPM data were refined by log2 fold change values (logFC) of PINK1 KO hDANs compared to control hDANs. We used Ingenuity Pathways Analysis (IPA, QIAGEN). First, twenty genes with the highest and lowest log2 fold change were listed. Based on terms ‘Parkinson’s disease’, ‘dopamine metabolism’ and known PINK1 interactions, several hit genes were validated in three independent differentiations by qRT-PCR. of sequencing data have been validated via quantitative reverse transcription-PCR (qRT-PCR). Interesting genes have been selected by the following criteria: a high logFC in sequencing sets, have a CMP-values >5 and occur in both PINK1 KO hDAN clones.

### Phosphate-affinity polyacrylamide gel electrophoresis (PhosTag SDS-PAGE)

For Parkin decorated Phos-Tag acrylamide gels, 100 µmol Phos-Tag (WAKO Chemicals) and 200 µmol MnCl2 were added to the SDS-PAGE and EDTA-free Laemmli buffer was used for sample loading. For all other experiemnts gels were prepared using 7.5% (v/v) polyacrylamide gel with 25 µM Phos-tag and 50 µM MnCl2 or Zinc-pre-cast PhosTag gels. Note that the prestained molecular weight markers contain EDTA and therefore do not resolve at the exact kDa on PhosTag gels. All PhoTag gels were run at ∼30V or slower. Phosphorylation-dependent mobility shifts of some proteins are dramatically improved by lowering the current. All gels were gently agitated in transfer buffer supplemented with 1 mM EDTA for 10 min then washed for 10 minutes in the transfer buffer alone before transfer. We used iBlot semi-dry blotting with nitrocellulose. Blocking, incubation with antibodies, and detection are according to standard Western blotting procedures.

### Phosphoproteomics

#### Samples preparation for mass spectrometry

Proteins from crude mitochondrial enrichment lysates were precipitated using acetone precipitation as described in (“Maximizing recovery of water-soluble proteins through acetone precipitation”; Crowell, Wall and Doucette; Analytic Chimica Acta 2013). Acetone precipitated proteins were denatured in urea buffer (6 M urea, 2 M thiourea in 10 mM Tris buffer, pH 8.0) and protein concentration was determined by applying Bradford assay (Bio-Rad). Disulfide bonds were reduced by 10 mM of dithiothreitol (DTT) for 1 h at RT under agitation at 800 rpm. In a next step alkylation with 50 mM iodoacetamide (IAA) was performed for 1 h at RT under agitation in the dark. Subsequent protein digestion in a ratio of 1 µg enzyme per 100 µg protein for 3 h with the endoproteinase Lys-C, and overnight with trypsin was performed. To stop the enzymatic reaction and to load peptides onto C18-material for purification, samples were acidified to pH 2.5 with trifluoroacetic acid (TFA).

For quantitative analysis samples were on-column (SepPak C18, Waters) labeled with dimethyl as described previously (Boersema, Nature Protocols volume 4, pages 484–494 (2009)). In more detail, two triple dimethyl sets were prepared, combining either the untreated or the simvastatin treated samples. Primary amines of peptides derived from WT were labeled with the light, KO1 with the medium and KO2 with the heavy labeled version of formaldehyde. Label efficiency of each dimethyl channel and the correct 1:1:1 mixing of each triple dimethyl set was validated by a pilot LC-MS/MS measurement. All label efficiencies were above 99% and the mixing correction factors for the two dimethyl sets were applied based on the label mixing ratios. Labeled peptides were eluted from the column by 80% acetonitrile (ACN). This left around 1.5 mg of labeled peptide mixture for the enrichment of phosphorylated peptides and 20 µg for the analysis of the proteome. Proteomic samples were subjected to purification by C18 StageTips as described previously (Ishihama, J. Proteome Res.200654988-994). Three consecutive rounds of phosphopeptide enrichment were performed by using MagReSyn®Ti-IMAC beads (ReSynBiosiences) according to the manufactures instruction. For all enrichment rounds 1 mg magnetic Ti-IMAC beads per 500 µg protein digest were used. Eluted peptides were purified by C18 StageTips prior to LC-MS/MS based analysis.

#### Mass spectrometry

Proteome and Phosphoproteome samples were measured on a Q Exactive™ HF Hybrid Quadrupol-Orbitrap instrument, online connected to an Easy1200 nanoflow HPLC system (Thermo Fischer Scientific).

The liquid chromatography-Mass spectrometry (LC-MS) analysis was performed as described in Schmitt et al https://doi.org/10.1074/mcp.RA119.001302. Briefly, the proteome samples were separated in a 230 min gradient of the binary solvent system compromising of solvent A and B on an in-house packed 20 cm reverse-phase C18 (1.9 µm; Dr. Maisch) nano-HPLC column. The phopsproteome samples were run in a 60 min gradient. The following instrument settings were used for the quantitative phosphoproteome analysis: full MS scans were recorded with a resolution of 60,000. The 7 most intense ions were selected for higher-energy collisional dissociation (HCD) fragmentation and ions were recorded with a resolution of 60,000. For the proteome analysis the 12 most intense peptides were selected and analyzed with a resolution of 30,000.

MS raw data were, processed by using Andromeda integrated in MaxQuant 1.5.5.1 (Cox J. Proteome Res.20111041794-1805; Cox Nat Protoc. 2009; 4(5):698-705) with default settings. Peak lists were searched against the target decoy UniProt human database (release 2019_02_13; 74,468 proteins. For quantification, the dimethylation on lysine residues and N termini was defined as “light labels” (+ 28 Da), “medium labels” (+ 32 Da), and “heavy labels” (+ 36 Da). The match-between-runs option between consecutive phosophoproteomic data was enabled. Oxidation of methionine and phosphorylation of serine (S), threonine (T) and tyrosine (Y) were set as variable modifications.

For a more accurate quantification the dimethyl ratios of identified phosphorylation events were normalized to the ratios of the corresponding protein groups by using Perseus software (version 1.5.3.2). Significantly regulated phosphorylation events log2-transformed dimethyl ratios were computed by the Significance B test and a p-value of 0.05.

### Crude mitochondrial enrichment

Crude mitochondrial enrichment: Fresh cell pellets on ice were resuspended in mitochondrial isolation buffer (10 mM HEPES pH 7.4, 50 mM sucrose, 0.4 M mannitol, 10 mM KCl, 1 mM EGTA, phosphatase and protease inhibitors (Roche)) and passed through 20G, 27G and 30G needles 8 times to disrupt cells. Fractionation was then achieved by several centrifugation steps. Fist, samples were centrifuged for 5 min at 1000 xg at 4°C and the supernatant was saved. The pellet was resuspended in 500 µL mitochondrial isolation buffer and passed through a 30G needle 8 times before centrifuging 5 min at 1000 xg at 4°C. The second supernatant was pooled with the first and centrifuged for 15 min at 15,000 xg at 4°C. The obtained pellet comprises the mitochondrial fraction and was resuspended in 300 µL PBS containing 1% TritonX-100, phosphatase and protease inhibitors. Mitochondria were lysed for 15 min on ice with repetitive vortexing.

### Cholesterol Assay

The Total Cholesterol Assay Kit (Fluorometric) (STA-390, Cell Biolabs Inc) was performed according to manufacturer’s instructions. Absolute cholesterol levels were measured using cholesterol standards and normalized to protein input in the lipid extraction.

### Lipidomics

Lipidomics of whole hDANs from independent differentiations (n=3) were performed according to the exact method described in ^99^.

Lipidomics of crude mitochondrial preparations from hDANs from independent differentiations (n=3) were performed according to the methods described in^100, 101^.

### Sucrose fractionation of hDANs and depletion of cholesterol

Detergent solubilisation of hDANs and subsequent sucrose gradient fractionation was performed as described in ^92^. The 14 fractions were stored at −80 °C and then proteins were precipitated using the methanol-chloroform method for all fractions to concentrate the sample sufficiently to detect proteins in the higher fractions.

### BRET Assay Mito-ER Tethering

All constructs for expression using Sindbis virus, were synthetized in pSinRep5 (Thermo Fisher Scientific). Restriction enzymes Mlu1 and StuI were used for the insertion of the constructs into the SR5 vector. Sindbis virus containing BRET biosensor (Donor: mitochondria targeted Renilla luciferase, Rluc and Acceptor: ER-targeted mVenus) were produced following protocol described in^102^, with minor modifications. Briefly, BHK-21 cells were co-transfected with pSinRep5 RNA of interest and helper pDHtRNA. After 48 hours, viruses were purified by sucrose gradient centrifugation. Samples were centrifuged 90 minutes at 35.000 rpm (4 °C) in a SW 60 Ti swinging-bucket rotor (Beckman Coulter) in a Beckman Optima L-100K. Viral particles were collected from the 20 %/55 % sucrose interface. NPCs and hDANS were infected with sindbis virus containing BRET biosensor for 48h. Cells were washed with PBS, incubated with 5 µM Coelentrazine h (Promega) in PBS for 10min in the dark. BRET measurements were carried out in a Tecan Infinite M200 plate reader at RT. BRET signal was calculated as acceptor emission relative to donor emission and corrected by subtracting the background ratio value detected when Rluc is expressed alone.

### Dynamic light scattering (DLS)

Dynamic light scattering is a technique used in physics for size characterization of proteins, nanoparticles, polymers and colloidal dispersions. hDANs were washed in PBS and suspended in 350µl of a standard HPLC elution buffer that does not contain detergents (see HPLC analysis). The hDAN suspension were homgenised using 5mm stainless steel beads and the tissue lyser LT (Qiagen) for 4 minutes with 50Hz.To remove large impurities the homogenized cell suspension was filtered through a 0.2µm nylon membrane. The filtered suspension was then diluted in water and run through a Malvern DLS instrument at 21°C. Particle size (nm) and sample polydispersity index (Pdi) are calculated automatically and analyses were made according to the manufacturer’s instructions. Samples with Pdi values close to 1 or 1 are generally considered too heterogeneous for the calculation of particle size and therefore these measuremenet were removed from the mean size data but not the mean Pdi data. *n*=diff 3.

### Neurotransmitter Uptake Assay

Neurotransmitter transporter activity in hDANs was measured using the Neurotransmitter Transporter Uptake Assay Kit (Molecular Devices) according to the manufacturer’s instructions. Mature hDANs were seeded in Matrigel coated black, clear bottom 96-well plates prior to the assay at a density of 60,000/well in triplicates. Wells containing no cells were used as an internal control. hDANs were treated with a specific VMAT2 inhibitor (Tetrabenazine, 10µM), a TH inhibitor (3-Iondol-L-Tyrosine, 300µM), or a combination of MAO and COMT inhibitors (Tranylcypromine 10µM and Tolcapone 100nM respectively). These compounds were added in media containing L-DOPA (50µM) or without L-DOPA. A subset of hDANs triplicates were treated with media alone or media containing L-DOPA (50µM). All treatments were incubated on the cells at 37°C for 20 minutes prior to the addition of the substrates. Uptake fluorescence was measured using the SpectraMax M_2_^e^ microplate reader in kinetic mode (Molecular Devices) measuring every 30s and using SoftMaxPro 6.4 Software (Molecular Devices) for detection. After the assay, the hDANs were washed and fixed in 4% (v/v) PFA containing Hoescht to account for cell number in each well and then counterstained for TH to account for any large differences in TH positive hDANs in the culture wells. Statistical analyses were accomplished by the unpaired two-sided Student’s t-test (*p<0.05, **p<0.01).

### Dopamine (Catecholamine) oxidation assay

The catecholamine oxidation assay was performed according to ^103^ using Biodyne® B 0.45µm membranes (Pall corporation) and detection using a LiCOR fluorescent scanning device.

### Monoamine oxidase (MAO) Activity Assay

MAO activity was monitored using a radiometric assay with 14C-tyramine hydrochloride as substrate as previously described^104^. Data were normalized for protein content and rates expressed as disintegrations of 14C/min/μg protein.

### Electrophysiology

Whole-cell patch clamp experiments were performed at room temperature using an Axopatch 200B amplifier (Molecular devices). Data were low-pass filtered at 10 kHz, digitized at 10 kHz via a Digidata 1440A acquisition system and analyzed offline using Clampfit 10.7 (Molecular Devices, Sunnyvale, CA, USA). During electrophysiological recordings, cells were kept in an extracellular solution (5mM HEPES (pH 7.4), 140 mM NaCl, 4.2 mM KCl, 1 mM MgSO4.7H2O, 1.1 mM CaCl2.2H2O, 0.5 mM Na2HPO4, 0.45 mM NaH2PO4, 10 mM glucose). Patch pipettes were pulled from borosilicate glass (Science products) using a Sutter P97 Puller (Sutter Instruments Company). Their resistance ranged from 3 to 5 MΩ. Patch pipettes were filled with intracellular solution (10 mM HEPES (pH 7.2), 5 mM EGTA, 135 mM K-gluconate, 4 mM NaCl, 0.5 mM CaCl2, 2 mM ATP-Mg, 0.4 mM GTP-Na) and an osmolarity of 290 mOsm. Series resistance (<20 MΩ) was monitored during the experiment. Cells showing unstable series resistance or resting membrane potential were discarded. Liquid junction potential was not subtracted from the data and it was calculated as −15 mV. The data are shown with leak subtraction. Voltage-clamp experiments were performed to record inward and outward currents generated via depolarization pulses from a holding potential of −70 mV to different potential levels, from −60 mV to +30 mV with 10 mV increments, for 24 milliseconds. To assess intrinsic firing properties, current-clamp experiments were performed holding the neurons at 55 mV level via continuous current injection and responses to the stepwise current stimulations from −0.001 nA to 0.069 nA in 0.005 increments with a duration of 800 ms were recorded.

### Calcium imaging

Mature neurons were seeded on Matrigel-coated glass coverslips. Fluo-4 Direct Calcium Reagent (Invitrogen) was added to the cells and incubated for one hour at 37°C and 5% CO_2_. After washing the cells with growth medium, the coverslips were transferred into the imaging chambers and Fluo-4 reagent diluted in growth medium was added. 3 mM EGTA was added per well 10 minutes prior to imaging. Neurons were imaged on a LSM 510 Confocal microscope (Zeiss) taking a picture every 0.5 sec for 25 minutes. A baseline was recorded for two minutes and then 2 µM Thapsigargin was added. The data analysis was performed using ImageJ. The corrected total cell fluorescence per cell was determined and plotted over time to evaluate the calcium content.

### Protein estimation

The protein content of cell supernatants was determined with a Pierce™ BCA Protein Assay Kit (ThermoFisher) and normalized to 20-80µg protein per lane depending on abundance.

### Mitochondrial Morphology (BacMam Mitogreen)

Neurons were plated on Matrigel covered glass coverslips, treated with CellLight™ Mitochondria-GFP, BacMam 2.0 (Invitrogen) according to manufacturer’s instructions for 24 hours in N2 medium. Mitochondria of transfected neurons were imaged using an LSM-510 confocal microscope (Zeiss). Z-stack pictures were analysed with ImageJ, to determine the aspect ratio, form factor, area and circularity of mitochondria according to ^105^.

### Kinetic analysis of mitochondrial membrane potential

Neurons on Matrigel covered glass coverslips were treated with N2 medium 24 h prior to the experiment. Kinetic TMRE measurements were conducted as described^106^ and imaged with a LSM-510 confocal microscope (Zeiss). The corrected total cell fluorescence was determined using ImageJ.

### Analysis of cellular ROS

Medium was changed to N2 medium 24 h prior to the experiment. On the following day neurons were incubated with N2 medium or N2 medium containing 10µM Rotenone for 4h before measurement. 100µM dihydroethidium (DiHET) (Santa Cruz Biotechnology) was added to all wells and fluorescence emission at 420nm (cytosolic DiHET, non-oxidized) and 610nm (chromatin-bound, oxidized DiHET) was measured every 30 secs for 30 min on a SpectraMax® M microplate reader (Molecular Devices).

### GSH assessment

Neurons on a 96-well plate were treated with different treatment conditions and N2 medium 24 h prior to the experiment. Cells were briefly washed with HBSS (Gibco) and incubated with 100µL HBSS containing 50µM Monochlorobimane (Sigma) for 40 minutes @37°C, 5% CO2. After washing with HBSS, cells were imaged with a SpectraMax® M microplate reader (Molecular Devices) (Ex 390nm/Em 478nm).

### Aconitase activity

Aconitase activity was measured in whole cell lysates from neurons using an adapted protocol^107^. Absorbance at 340 nm was measured using a SpectraMax® M microplate reader (Molecular Devices).

### NAD^+^ and NADH levels

Neurons were cultivated in N2 medium 24 h before the experiment. Whole cell NAD^+^- and NADH-levels were determined using the NAD/NADH Assay Kit (Fluorometric) from Abcam according to manufacturer’s instructions. NADH reaction mixture was incubated for 1 hour and fluorescence (Ex/ Em = 540/ 590 nm) was measured using a SpectraMax® M microplate reader (Molecular Devices).

### Complex I and citrate synthase activity

Complex I activity was measured in crude mitochondrial enrichments from hDANs previously described in ^105^. Following isolation of crude mitochondria, a total protein content >0.7mg/mL is required to have enough active mitochondria for each sample in triplicate in the assay and again duplicated for the rotenone negative control. Complex I activity was measured according to^108^. The activity of complex I was normalized to citrate synthase activity, also according to ^108^ and data expressed as a ratio of complex I/citrate synthase.

### Transmission Electron microscopy

Neurons were seeded on Matrigel (Corning) coated glass coverslips and cultivated for three days prior to treatment with N2 medium with and without 1 µM Valinomycin for 24 hours. After washing and fixation with 2.5% glutaraldehyde (Science Services, Munich, Germany) in cacodylate buffer (pH7.4; Merck-Millipore, Darmstadt, Germany) overnight at 4°C, cells were washed with cacodylate buffer, post fixed in 1% osmiumtetroxide, dehydrated and embedded in epoxide resin (Araldite, Serva, Heidelberg, Germany) as described previously ^109^. Ultrathin sections were performed using a Reichert Ultracut ultramicrotome (Leica, Bensheim, Germany) and were analyzed in an EM 10 electron microscope (Zeiss, Oberkochen, Germany). Images were taken by a digital camera (Tröndle, Germany).

### Measurement of biogenic amines by HPLC-ED

Mature hDANs were expanded during differentiation into 175cm^2^ Matrigel coated flasks. For each experiment, approximately 2-3 175cm^2^ flasks were required. Although endogenous dopamine levels could be detected in hDANs derived from much larger cell volumes, we employed 50µM L-DOPA treatment overnight (16h) to enhance dopamine metabolism without risk of dopamine toxicity (see ^103^ and ^110^). Following detachment of hDANs with Accumax, the cell suspensions were washed in PBS and cells counted. PBS was used to normalise the number of cells in the suspension. An aliquot of the normalized suspension was taken for preparation of protein lysate and RNA to determine the relative amount of TH positive cells in each experiment and differentiation for each cell line. Fresh (unfrozen) cell pellets kept on ice were suspended in 350µl of a standard HPLC elution buffer that does not contain detergents (Thermo Scientific). The suspensions were homogenized using 5mm stainless steel beads and the tissue lyser LT (Qiagen) for 4 minutes with 50Hz.To remove large impurities the homogenized cell suspension was filtered through a 0.2µm nylon membrane. Samples were analysed for catecholamine and indolamine content by ion-pair reverse phase HPLC with coulometric detection (Ultimate 3000 LC with electrochemical detection ECD3000RS, Thermo Fischer Scientific, California, USA). A hypersil C18 colum was used (150×3 mm, 3 µm) and the system was run with a Test mobild phase containing 10% actonitril and 1% phosphate buffer (Thermo Fischer Scientific, California, USA) at a flow rate of 0.4 ml/min at 30 °C. The potential of the first channel was set to +350 mV, the second channel to −250 mV.

Epinephrine, Norepinephrine, Dopamine, 3,4-dihydroxyphenylacetic acid (DOPAC), homovanillic acid (HVA), 5-hydroxyindol-acetic acid (HIAA), 3-Methoxythyramine (3-MT) and serotonin (5-HT), concentration was determined by comparing peak areas of the samples with those of standards using Chromeleon 7 chromatography data system software. The neurochemicals in standards were determined with a high correlation linearity (r2 = 0.98) and good reproducibility in retention time (0.03%). The limit of detection was <1 pg on column for all the metabolites analyzed.

### Complex I Dipstick Assay

Active Complex I was pulled down from hDAN homogenates using the Complex I Dipstick Assay from AbCam (ab109720) according to the manufacturer’s instructions.

### Mitochondrial Import Assay

Radiolabeled proteins were synthesized in rabbit reticulocyte lysate in the presence of 35S-methionine after in vitro transcription by SP6 polymerase from pGEM4 vectors (Promega). Radiolabeled precursor proteins were incubated at either 30°C (pSu9-DHFR) or 4°C (Fis1) in import buffer (250 mM sucrose, 0.25 mg/ml BSA, 80 mM KC1, 5 mM MgCl2, 10 mM MOPS-KOH, 2 mM NADH, 4 mM ATP, pH 7.2) with mitochondria isolated from the indicated cells. Non-imported pSu9-DHFR was removed by treatment with proteinase K (PK, 50 µg/ml) for 30 min on ice and then PK was inhibited with 5 mM PMSF. Membrane integration of Fis1 molecules was confirmed by resistance to alkaline extraction (incubation on ice for 30 min with 0.1 M Na2CO3). Finally, all the samples were boiled at 95°C for few min before their analysis by SDS-PAGE.

## Acknowledgements

The German Research Foundation and the research training group MOMbrane and The German Federal Ministry of Education (BMBF) ‘Mito PD’ were the primary source of funding and allowed collaborations that made this work possible. Funded by the German Research Foundation (DFG), GRK2364.

Maria Calleja-Felipe of the Molecular Cognition Laboratory, Biophysics Institute, Consejo Superior de Investigaciones Cientificas (CSIC)-University of the Basque Country (UPV)/Euskal Herriko University (EHU), Campus Universidad del País Vasco, 48940 Leioa, Spain.and, Shira Knafo of the Molecular Cognition Laboratory, Biophysics Institute, Consejo Superior de Investigaciones Cientificas (CSIC)- University of the Basque Country (UPV)/Euskal Herriko University (EHU), Campus Universidad del País Vasco, 48940 Leioa, Spain. Ikerbasque, Basque Foundation for Science, 48013 Bilbao, Spain. Department of Physiology and Cell Biology and National Institute of Biotechnology in the Negev, Faculty of Health Sciences, Ben-Gurion University of the Negev, Beer-Sheva, 8410501 Israel who developed the constructs for performing the BRET experiments for measurement of mitochondrial-ER contacts and kindly shared them with HFR and AG. Gerrit Machetanz of the Hertie Institute for Clinical Brain Research, The University of Tübingen, University Clinic Tübingen for clinical assistance and discussions concerning PINK1 Parkinson’s disease.

## Author contributions

1. **Research project: A.** Conception (JCF, TG, PK), **B**. Organization (JCF, SG, RK, TG, PK, AG, BM, HK, DVW, WW), **C**. Execution (JCF, CB, BB, NC, PFB, MF, HFR, SG, AS, BS, LS, AUK, BU, MZ, KZ, DVW).
2. **Statistical Analysis: A**. Design (JCF, CB, AS, MF, NC, PFB, SG, KZ, BM, BB, HFR, HK, DR) **B.** Execution (JCF, CB, AS, MF, NC, PFB, SG, KZ, BM, HFR, HK) **C**. Review and Critique; (JCF, SG, PK, TG, MF, AS, NC, AG, HFR, HK, AUK, KZ, BM, DVW, WW, DR)
3. **Manuscript**: **A**. Writing of the first draft (JCF), **B.** Review and Critique (DVW, JCF, SG, RK, TG, AG, MF, HFR, PK, AUK, KZ, BM, WW).

## Conflict of Interest

The authors declare no conflict of interest.

## The paper explained

### Problem

Parkinson’s disease is a common neurodegenerative disease that is currently incurable. There is little known about why dopamine-containing neurons are so vulnerable. It is not clear how mitochondrial defects (that occur in all body cell types) cause neuronal dysfunction and neuronal death in Parkinson’s disease.

### Results

We studied dopaminergic neurons grown from healthy and Parkinson’s disease stem cells. Mitochondrial vulnerability induced by defective PINK1 (a Parkinson’s disease gene) is compensated for in maturing human neurons and mitochondrial networks are normal. A compensatory metabolic shift provides lipids for piecemeal renewal of mitochondrial membranes in the absence of PINK1-dependent mitophagy. However, these biochemical events have a negative impact at mitochondrial-ER contact sites (affecting central metabolism and dopamine metabolism) and at the plasma membranes making them more ridged and therefore more susceptible to synaptic defects (including dopamine uptake).

### Impact

These data highlight the complexity and difficulties in treating Parkinson’s disease. Here the biological mechanism uncovered is both necessary for the survival of neurons during development but leads to their vulnerability in aging. We propose that low doses of compounds such as β-cyclodextrin which make cell membranes more fluid could benefit patients by treating the secondary (but more specific) cause of disease rather than the initial defect which has so far proven unsuccessful.

## Data availability

- RNA Seq data: All raw data and log2 values for the genome.
- Phospho-proteome data: All raw data and phosphorylation sites of all unfiltered and filtered data.
- Lipidomics: All lipidomics data.

Links to Excel and Text files will be made via https://datadryad.org/stash and a link via https://www.hih-tuebingen.de/en/forschung/neurodegenerative-diseases/research-groups/mitochondrial-biology-of-parkinsons-disease/

The data will be uploaded on datadryad if this manuscript is accepted for peer review so it is accessible for reviewers.

**Supplementary Figure S1A)** Relative PINK1 gene expression in control and PINK1 KO hDAN clones using primers directed to exon1-4 and exon 2-4 (n=diff3). **S1B)** Immunocytochemistry staining of MAP2, TH and FOXA2 in fixed hDANs. Representative images are shown (n=diff4). **S1C)** Representative images of CI dipstick assay for active CI in hDANs treated with or without 1µM valinomycin for 24h (left panel). Quantification of dipstick band from three independent experiments (n=diff3). **S1D)** Mean average TMRE fluorescence signal during flow cytometry for untreated hDANs and those treated with 10µM CCCP in vitro. The signal is normalised to the unstained signal from the same cells. n=diff 3. **S1E)** Kinetic measurement of dihydroethidium-cytosolic ROS over time in the presence or absence of 1mM rotenone. **S1F)** Lipid peroxidation levels measured using BODIPY in hDANs treated with rotenone, L-DOPA, DA and 6-hydroxydopamine (6-ODHA) for 24h. **S1G)** Glutathione assay in hDANs in the presence or absence of rotenone, glutathione depleting BSO or both together. n=diff3. **S1H)** alpha-ketoglutarate dehydrogenase enzyme activity in hDANs. n=diff3. **S1I)** Total NADH levels in hDANs. n=diff3.

**Supplementary Figure 2A)** Heatmap of strongest log2 fold changes in gene expression between control and PINK1 KO genotypes in hDANS (n=3, diff=1). **S2B)**% mol lipid of all measured lipid species from lipidomics of mitochondrial fractions of hDANs (n=diff3). **S2C)** % mol lipid of all measured lipid species from lipidomics of whole hDAN preparations (n=diff3). **S2D)** Co-localisation (Pearson coefficient) of SREB to nuclei in fixed hDANs (n=3). **S2E)** Readouts of mitochondrial morphology in hDANs from live cell imaging (n=diff3). **S2F)** Phosphorylation status of TOM70, Mitoferrin2 (MFRN2) and GRP75 in NPC neuroprogenitors and hDANs where CCCP is used to depolarise the MOM and stabilise PINK1 (representative blots, n=diff3). **S2G)** List of most regulated (log2 fold change) phospho-peptide abundance in hDANs untreated or treated with 5µM Simvastatin for 24h. The list was filtered and weighted for the lowest deviation between samples and *represents the fold change is in the same direction and **reversed by Simvastatin. **S2H)** Phosphorylation status of GAP43 and PDRMC1 in hDANs where CCCP is used to depolarise the MOM and stabilise PINK1 (representative blots, n=diff3).

**Supplementary Figure 3A)** Relative Rab3B gene expression and **3B)** Relative PNPO gene expression in NPC neuroprogenitors and hDANs (n=diff 3). **3C)** Cholesterol levels measured in the ventral midbrain (VM, left) and striatum (STR, right) of WT (CTRL) and PINK1 KO (KO) mice at 4 months (young) or 16 months (old) of age. n= 3 animals per group. Mean average cholesterol +/− standard deviation. *=<0.05, Student’s T-test. **3D)** Representative SDS PAGE Western blot images of mitochondrial protein import for pSU9-DHFR and Fis-1 (left) and quantification of SDS-PAGE Western blot images from four independent differentiations and experiments on fresh mitochondria (right).

